# Resource supplementation alters host exposure, susceptibility, and infection dynamics across a diverse parasite community

**DOI:** 10.64898/2025.12.17.694884

**Authors:** Amy R Sweeny, Gregory F Albery, Saudamini Venkatesan, Agata Delnicka, Andy Fenton, Simon A. Babayan, Amy B. Pedersen

## Abstract

Resource availability is crucial in shaping exposure and resistance to parasites. Although effects are expected to vary across parasite species, most of our knowledge comes from single-host, single-parasite studies. In a wild wood mouse population (*Apodemus sylvaticus*), we supplemented dietary resources over 3 years and quantified effects on a diverse parasite community alongside host behaviour and physiology. Resource supplementation had diverse impacts on 12 parasite taxa including positive (3/12), negative (6/12), and null (3/12) results. Supplementation also influenced host biology, with supplemented mice moving less, experiencing higher conspecific densities, improved conditions, and increased reproduction. These findings experimentally demonstrate that resource availability can multifacetedly affect disease dynamics. The complexity likely arises due to combined effects on exposure and susceptibility processes. Our study advances understanding of disease drivers at the human-wild interface and underscores the importance of assessing whole parasite communities to predict responses to changing resource landscapes in the wild.

## Introduction

Resource availability plays a crucial role in determining both individual- and population-level dynamics of infectious diseases (Bogdziewicz & Szymkowiak 2016; Calder & Jackson 2000; Randolph & Storey 1999). Increasingly, anthropogenic activities are altering the quality and quantity of resources available to wildlife, either via intentional management efforts or incidental exposure to human waste (Oro *et al*. 2013). Such supplemental resources can significantly alter host condition, immune investment, demography and behaviour, pathways with strong potential to reshape infection dynamics (Becker *et al*. 2015). While improved nutrition may enhance infection resistance via increased resources allocated to immune function (Calder & Jackson 2000), outcomes are context-dependent and can be neutral or even detrimental (Becker & Hall 2014), particularly if supplemental food is of poor quality, as shown in coyotes consuming urban waste (Murray *et al*. 2016). The outcome of resource supplementation for infectious disease will therefore depend on the quality of food source, and on the myriad ways altered resources may impact the host (Becker *et al*. 2015; Erazo *et al*. 2022, 2025).

The consequences of resource supplementation for disease outcomes arise from effects on both within-host processes, which alter susceptibility, and between-host processes, which modify exposure to parasites Exposure can increase via aggregation around food sources and changes in movement, population density, and social contacts with infected conspecifics (Mistrick *et al*. 2024). An increasing number of theoretical and empirical studies have demonstrated effects of resource availability are dominated by host demography and spatio-social behaviour. For example, a recent experiment that increased the density of bird feeders resulted in significantly increased *Mycoplasma gallisepticum* transmission within house finches (*Haemorhous mexicanus*) (Moyers *et al*. 2018), and intentional resource provisioning among Rocky Mountain Elk (*Cervus canadensis*) populations resulted in increased infection with *Brucella abortis* (Cross *et al*. 2007).

However, resource-driven changes in susceptibility and exposure are unlikely to act independently, or uniformly across individuals and parasites. Alongside effects on immunity, improved condition may increase reproductive effort and activity, elevating contact rates, or alternatively reduce foraging requirements lower exposure risk (Glaudas & Alexander 2017; Nagy & Holmes 2005). This context dependence is further shaped by parasite traits, where varied virulence strategies will shape the relationship between resources and immunity within-host (Cressler *et al*. 2014), and transmission mode is likely to influence whether changes to exposure or susceptibility have a stronger impact on infection outcome. A meta-analysis of 132 records confirmed varied direction and magnitude of resource supplementation effects across parasite taxonomic groups (Becker *et al*. 2015), yet how these interacting pathways manifest to shape infection outcomes within a single wild population containing multiple co-circulating parasites remains poorly understood.

Co-infection is ubiquitous in wild populations, and mammals typically host diverse parasite assemblages throughout their life (Cox 2001; Petney & Andrews 1998). Resource-driven physiological and behavioural changes are expected to act simultaneously across the entire parasite community, making the net outcome difficult to predict. However, empirical tests of these effects remain incredibly rare, largely because they typically require experimental manipulation and repeated screening across full parasite communities (Hellard *et al*. 2015; Pedersen & Fenton 2015).

Here, we use a large-scale resource supplementation experiment in a naturally co-infected small mammal population to ask how resource supplementation affects host behaviour, condition, immunity, and infection outcomes across a diverse parasite community. Wild wood mice (*Apodemus sylvaticus*) are found throughout Europe and are commonly infected with a diverse and abundant community of parasites (Knowles *et al*. 2013). We have previously shown that experimental supplementation of resources increases resistance to, and lowers infection burdens of, the common gastrointestinal nematode *Heligmosomoides polygyrus*, while also positively impacting anthelmintic drug efficacy (Sweeny *et al*. 2021c). Here we investigate the impact of supplementation with a high-quality resource on the broader micro- and macroparasite community, including gastrointestinal parasites (helminths and protozoans), ectoparasites, and blood-borne parasites (protozoans, bacteria, and viruses). Additionally, we used morphometric information, trapping data, and samples collected at each capture to quantify host social and spatial behaviour, reproductive effort, immune responses and body condition.

We show that food supplementation significantly impacted the parasite community, with some parasite species decreasing in infection probability and intensity, while others remained unaffected or even increased. The experimental supplementation of resources, while only conducted over a relatively short period of time, also had substantial impacts on host behaviour, condition, and demography. By integrating behavioural, physiological, and infection outcomes, we provide a robust experimental test of how resource enrichment reshapes parasite community dynamics in a wildlife population. These results highlight the importance of understanding direct and indirect impacts of provisioning for predicting response to altered resource availability for wildlife populations.

## Methods

### Field Experiment

Trapping took place from 2015-2017 in Callendar Wood (55.990470, −3.766636; Falkirk, Scotland), a 100ha broadleaf woodland, which contains a population of wood mice which are naturally exposed to and infected with a wide range of parasites. The experiment had three temporal replicates; all of which took place during the wood mouse breeding season: (i) May - July 2015 (ii) June - August 2016 and (iii) July-November 2017. Nutrition was manipulated at the population level by supplementing resources (unit – trapping grid; control (unmanipulated) vs. supplemented nutrition). Within each grid, parasite communities of some individuals were perturbed with anthelmintic treatment at the individual level (control (water) vs. anthelminthic treatment). Full details of the methods and trapping grids can be found in (Sweeny *et al*. 2021c). Trapping duration was extended in 2017 to monitor the duration of any supplementation effects. Briefly, we carried out resource supplementation for three weeks prior to the beginning of the trapping and then continued throughout the sampling period. We supplemented grids twice per week with 2kg/ 1000m^2^ of sterilized, TransBreed™ mouse chow pellets, scattered at regular intervals across the grids to ensure an even spatial distribution. TransBreed™ is a high-nutrient, standard veterinary feed which is formulated for optimum breeding performance in laboratory mice and offers whole-diet nutrition to the wild mice in this study (20% protein, 10% fat, 38% starch, high content of micronutrients). Following this 3-week period of supplemented nutrition, we live-trapped mice for 3 nights/week using Sherman live traps (H.B. Sherman 2×2.5×6.5 inch folding trap, Tallahassee, FL, USA). In 2017 only, from mid-September grids were trapped every other instead of every week for the autumn season. Traps were baited with cotton wool bedding, seeds, carrot, mealworms, and TransBreed pellets (on supplemented nutrition grids only), set in the early evening (16.00-18.00) and then checked early the following morning. All wood mice weighing >10g were tagged with a subcutaneous microchip transponder for identification (Friend Chip, AVID2028, Norco, CA, USA). On both control and nutritional supplementation grids, mice at first capture were rotationally assigned within each sex to either control or anthelminthic treatment groups. We used a weight-adjusted of 2ml/g dose of both Pyrantel pamoate (Strongid-P, 100 mg/kg) and Ivermectin (Eqvalan, 9.4mg/kg). In 2017, treatment was re-administered 4 weeks after initial dose for all individuals still in the experiment.

Each tagged individual was followed for a period of 12-16 days (2015-2016) or up to 8 weeks (2017). During this time, we collected the following morphometric data at every capture: sex, age, measures of host condition, including body mass, length, fat scores, and reproductive status (as described in (Sweeny *et al*. 2021c)). Blood samples were collected via mandibular bleed (first capture, or first capture and one month later in 2017) or tail snip (subsequent captures) a maximum of once per week; faecal samples were collected for each mouse at every capture from previously sterilised traps and preserved in 10% formalin.

### Parasitology

Parasite abundance was estimated for GI helminths and coccidia as eggs or oocysts per gram (EPG or OPG) of faeces through salt flotation methods and microscopy taken from samples collected throughout the experiment as described in (Knowles *et al*. 2013). Briefly, eggs and oocysts were counted and standardized by the weight of the sample to estimate EPG and OPG; those values were rounded to the nearest integer for subsequent analysis. Ectoparasites (mites, fleas, and ticks) were counted for each individual following fur-brushing in the field. In 2015 and 2016, we also screened for three blood parasites (Wood Mouse herpes virus (WMHV), *Bartonella spp*. and *Trypanosoma grosi*) using diagnostic PCR. Blood samples collected in the field were returned to the laboratory and spun in a centrifuge at 12000 rpm for 10 minutes, and serum and blood pellet were separated and stored at −20° until processing. Details of blood DNA extraction and PCR diagnostics can be found in the supplementary material.

### Behaviour

We calculated spatial, social and density metrics to assess the impact of supplementation on host behaviour. Measures included: (i) **Density −** the density of individuals’ centroids in the sampling year, calculated from annual density kernels, (ii) **Max Distance −** The maximum Euclidean distance between known trapping locations for an individual across the study period, (iii) **Sum Movement −** The cumulative distance travelled between trapping locations by an individual for subsequent trapping sessions, (iv) **Strength −** the sum of the edge weights (total edges) for individual nodes within the network, where edges were defined as unique mice trapped nearby (within one adjacent trap distance) in the same trapping night, and (v) **New Births −** the number of juveniles recruited to a grid per trapping session. We computed density by creating space use distribution kernels with the adehabitathr package in R. We rasterized the usage distribution, assigning each individual a local density based on the raster value for their grid location.

### Condition and immunity

We quantified individual immune responses from at each capture from stored blood serum via a) the absolute concentration of total faecal IgA and b) relative, standardised concentration of *H. polygyrus*-specific IgG1 as reported previously (Sweeny *et al*. 2021c). Briefly, we calculated total faecal IgA concentration by extrapolation from a standard curve of known concentrations from a synthetically manufactured standard antibody. *H. polygyrus*-specific IgG1 was calculated as a relative concentration to a positive reference sample consisting of sera from *Mus musculus* experimentally infected with *H. polygyrus* in the laboratory. We refer to both IgA and IgG1 values as ‘antibody concentration’. Full details of immune assays can be found in the supplementary material.

Nutritional status was assessed by quantifying circulating serum albumin and total protein concentration from stored serum samples. Serum albumin is a dynamic and long-lived plasma protein which has important functions for multiple physiological roles (Garnier *et al*. 2017). Assays were optimized from (Garnier *et al*. 2017) for use on mouse samples using samples from our University of Edinburgh colony of formerly-wild wood mice. Full details of immune assays can be found in the supplementary material.

We assessed the condition of individuals at each capture using four metrics: i) body mass (g), ii) body fat score, iii) scaled mass index (SMI), and iv) condition (residuals of a weight by body length regression). Fat scores were assessed on a scale of 1-5 (emaciated-obese) by palpating the sacroiliac bones (back and pubic bones) as detailed in (Ullman-Cullere & Foltz 2011). SMI was calculated to provide a measure of condition scaled to allometry as detailed in (Peig & Green 2009).

### Statistical Analysis

All statistical analysis was run using R v 4.2.3 (R Core Team 2023). All models were run using the package ‘INLA’ (Rue *et al*. 2009).

We investigated the effect of resource supplementation on host behaviour using four GLMMs (one for each behaviour metric calculated). We fit each metric described above as the response variable with gaussian error families for scaled movement metrics, Poisson error families for network strength, and negative binomial error families for new births. Similarly, we investigated the effects of supplementation on host condition and immunity using eight GLMMs (one for each condition or immunity metric).

We assessed the impact of supplementation on parasite infection using 12 GLMMs (one for each parasite species/group). When the data were available for specific parasite taxa, we used abundance of infection (eggs/ oocysts per gram of faeces count or ectoparasite count) as the response variable with negative binomial error families. For the parasite species where abundance was not measured, or where distributions were too overdispersed to fit count models, we used probability of infection (presence/absence) as the response variable. For fleas, although we did collect count data, a very high percentage (99%) were 0 or 1 counts; so we just investigated the probability of infection (presence/absence) for this ectoparasite.

We fit the following fixed effects for each behaviour, condition and immunity metric, or parasite considered: supplementation (control or supplemented), year (2015, 2016, or 2017), sex (male or female), and reproductive status (active or inactive). The response ‘new births’ had an additional fixed effect of week (continuous) to account for growth over time. All probability or abundance of infection models had an additional fixed effect of scaled mass index (continuous), to investigate whether body condition facilitates infection with each parasite. Because randomised anthelmintic treatment was carried out for other experimental purposes during trapping, we also account for a fixed effect of drug treatment (control or treated) in each model.

Finally, we tested whether changes in behaviour or condition might mediate effects of experimental supplementation on the parasite community by testing additions of any behaviour or condition metrics significantly influenced by supplementation as fixed effects in GLMMs modelling infection as a response. If behaviours that are affected by supplementation are related to infection in this population, we expect that these behavioural shifts are likely to play a role in overall changes to the parasite community following supplementation.

## Results

### Parasite community of wild wood mice

Wood mice in our study populations harboured a diverse community of macro-, micro-, and ecto-parasites (Figure 1). The nematode *H. polygyrus* was the most common gastrointestinal (GI) parasite, followed by the protozoan *Eimeria hungaryensis*. Ticks were highly prevalent, with other ectoparasites detected infrequently. Blood-borne microparasites screened for in 2015-2016 were nearly as common as the GI parasites. For example, Gram-negative bacteria *Bartonella spp*. were detected in over 35% of mice.

**Figure 1.**
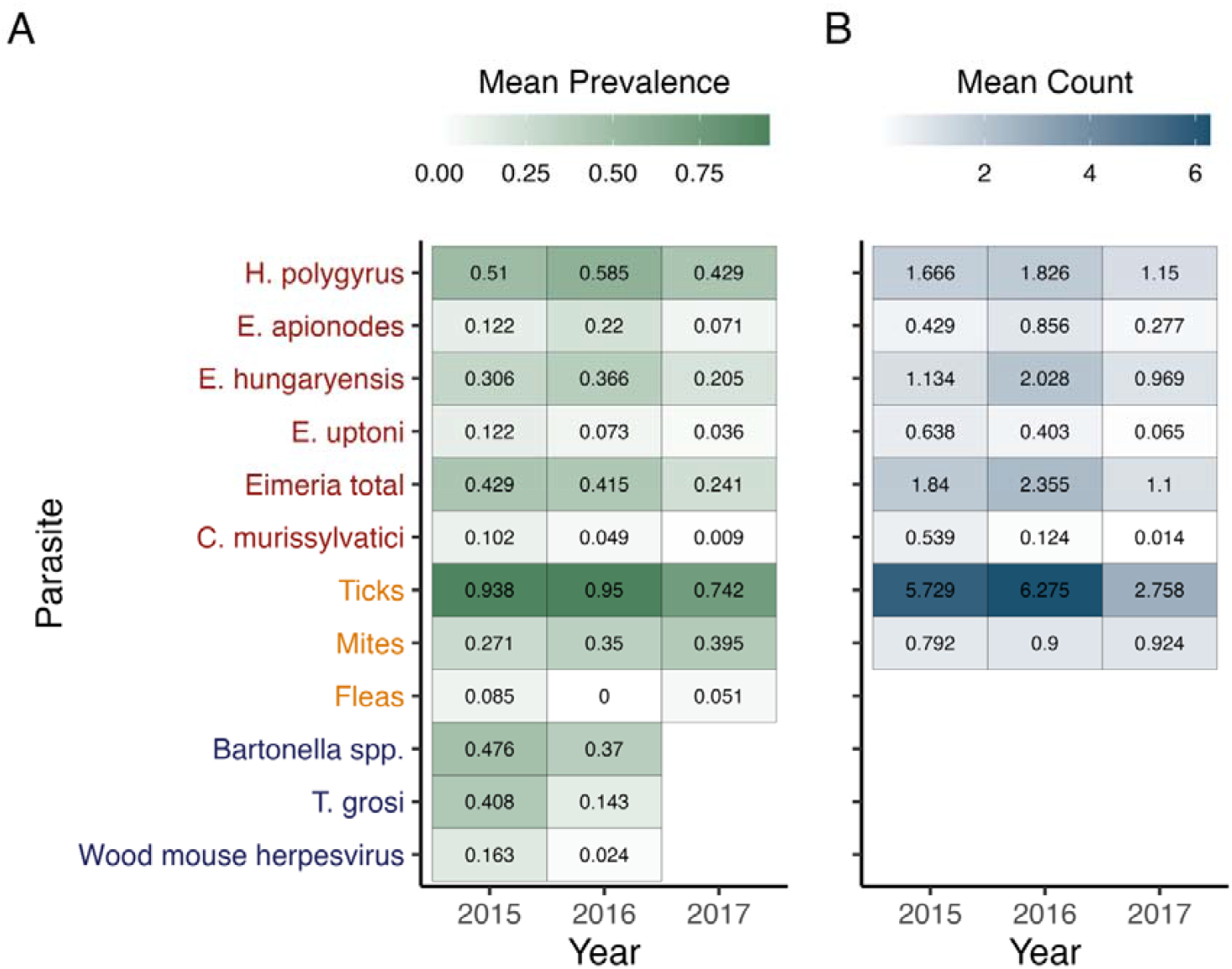
The parasite community of wild wood mice prior to food supplementation. A. Mean prevalence across years B. Mean count (mites, ticks) or log count (GI parasites). Mean count for gastrointestinal parasites (red) represents the mean log count + 1 egg or oocyst per gram faeces. Mean count for ectoparasites (yellow) represents the number counted visually. Values are calculated from first captures of all mice before treatment. Parasite labels: Red - gastrointestinal; Yellow - ectoparasite; Purple - bloodborne).

### Supplementation dramatically alters host behaviour & demography

Resource supplementation significantly affected the behaviour of mice on supplemented versus control grids for all metrics tested (Figure 2A). Mice on supplemented grids moved less than their control counterparts. Cumulative movement between sequential captures throughout the trapping season was lower for supplemented grids (Sum Movement: Estimate = −0.52, sd = 0.20, p = 0.010), as was the total euclidean distance covered by individuals across all trapping sessions (Max Distance: Estimate = −0.46, sd = 0.22, p = 0.033). Contact networks on supplemented grids were also less connected than on control grids, such that supplemented individuals had lower numbers of total contacts with other mice (Strength: Estimate = −0.44, sd = 0.15, p = 0.0047). In contrast, supplemented grids recruited more juveniles (New Births: Estimate = 0.87, sd = 0.14, p < 0.0001) and had higher densities (Density: Estimate = 0.84, sd = 0.11, p < 0.0001) than control grids. Aspects of behaviour were also influenced by factors such as sex, year, and reproductive status (Figure 3, Table S1).

**Figure 2.**
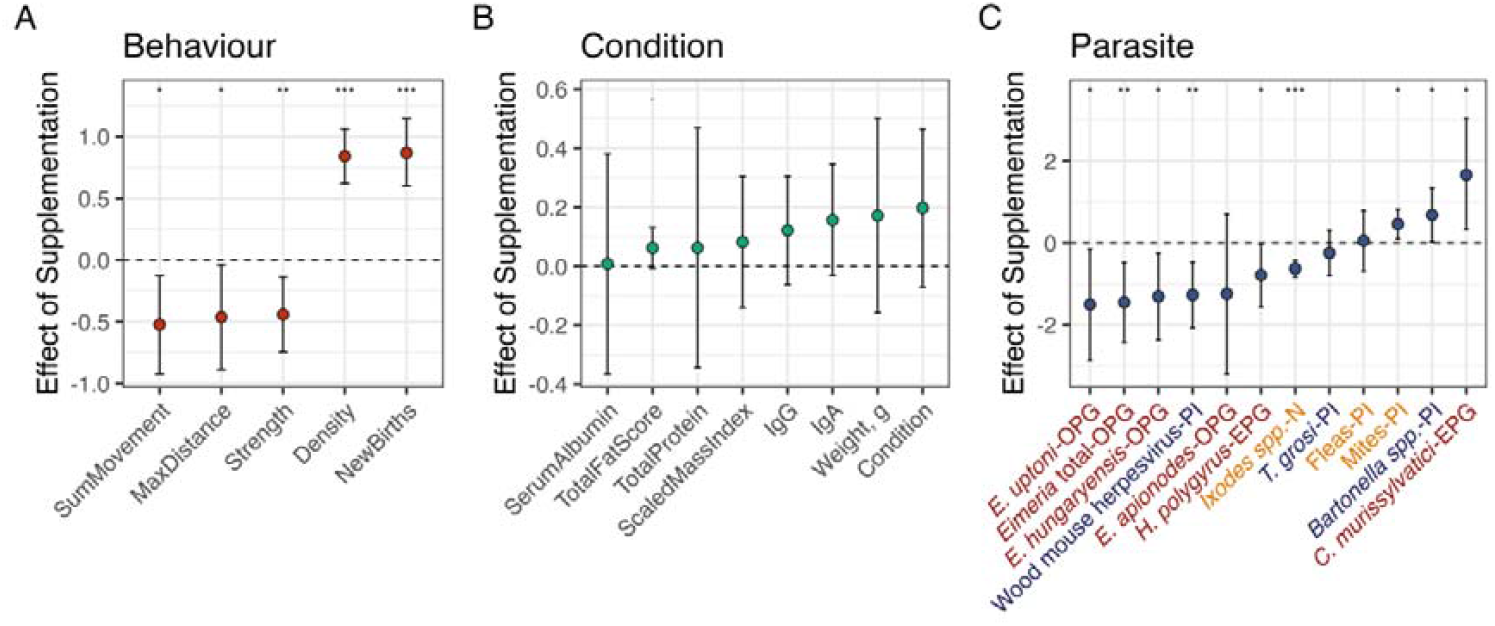
Estimates for the effect of resources (supplemented vs. control) on responses from model sets acrossA. Behaviour, B. Condition & Immunity, C. Parasites. Points and error bars respectively represent the mean and 95% credibility intervals for posterior distributions of effect estimates. Type of data modelled is indicated after parasite names (Counts = OPG, EPG, N; Probability of infection = PI). Parasite labels: Red - gastrointestinal; Yellow - ectoparasite; Purple - bloodborne).

**Figure 3.**
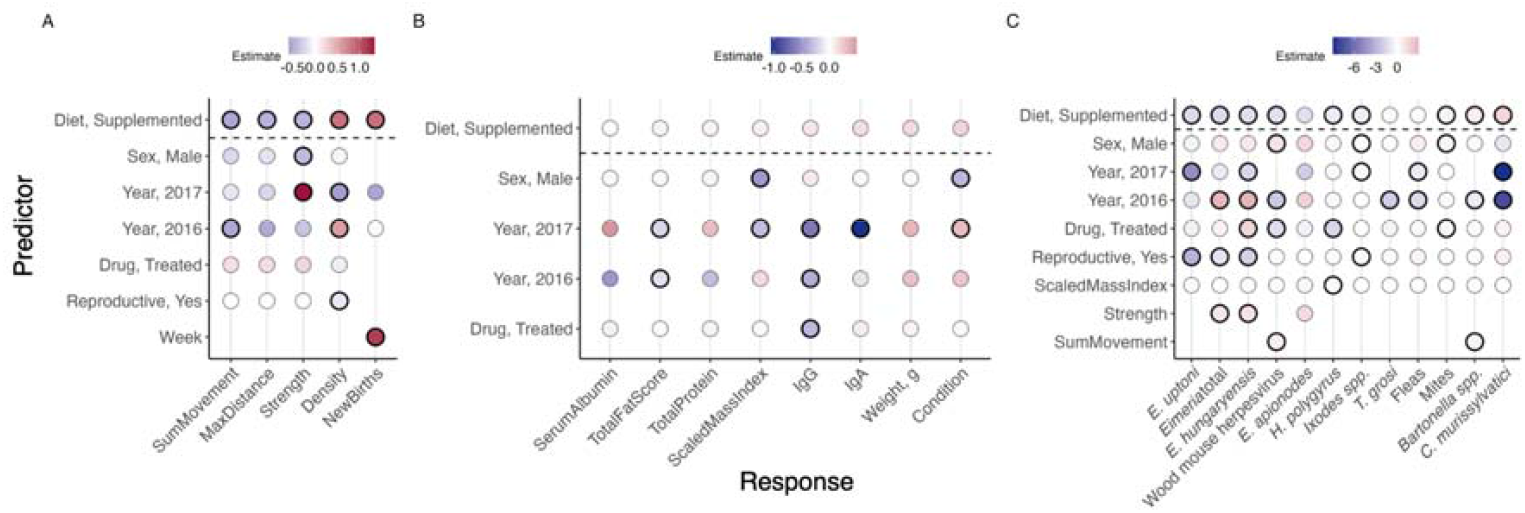
Full model effect estimates for A. Behaviour, B. Condition & Immunity, and C. Parasites. Shading represents effect estimates of each predictor (y-axis) for each response across model sets (x-axis). Significant results are indicated by the presence of a thick black border for reported estimates.

### Supplemented individuals have slightly improved condition

Effects of resource supplementation on metrics of nutritional status, condition and immunity levels varied considerably (Figure 2). Circulating serum albumin and total protein concentrations did not vary between supplemented and control mice, but there was a trend of increased condition and immunity measures among supplemented mice (Total Fat Score: Estimate = 0.062, sd = 0.035, p = 0.077; Scaled Mass Index: Estimate = 0.082, sd = 0.11, p = 0.47, Weight, g: Estimate: 0.17, sd = 0.17, p = 0.30; IgA: Estimate = 0.16, sd = 0.096, p = 0.10; IgG: Estimate = 0.12, sd = 0.093, p = 0.19). Sex and year also influenced condition measures, and year and drug treatment influenced antibody levels (Figure 3, Table S2).

### Resource supplementation has diverse effects across the parasite community

Resource supplementation had a broad significant impact across the parasite community, but not in a consistent direction (Figure 2). Mice on supplemented grids had significantly lower infection intensities (“OPG”/”EPG”/”N”) or probability of infection (“PI”) of *E. uptoni* (OPG, Estimate = - 1.50, sd = 0.69, p = 0.029), *E. hungaryensis* (OPG, Estimate = −1.31, sd = 0.54, p = 0.015), total *Eimeria spp* (OPG, Estimate = −1.45, sd = 0.49, p = 0.0034), wood mouse herpesvirus (PI, Estimate = −1.27, sd = 0.41, p = 0.002), *H. polygyrus* (EPG, Estimate = −0.78, sd = 0.39, p = 0.043), and *Ixodes spp*. (N, Estimate = −0.62, sd = 0.10, p < 0.001). Conversely, supplementation increased infections of mites (N, Estimate = 0.46, sd = 0.18, p = 0.013), *Bartonella spp*. (PI, Estimate = 0.68, sd = 0.33, p = 0.042), and *C. murissylatici* (EPG, Estimate = 1.66, sd = 0.69, p = 0.015). *E. apionodes, T. grosi*, and fleas were unaffected by supplementation. Abundance and prevalence across the parasite community was also influenced by a number of other factors including host sex and condition, year, and drug treatment (Figure 3, Table S3). For example, anthelmintic drug treatment decreased abundance of *H. polygyrus* (EPG, Estimate = −1.65, sd = 0.37, p < 0.0001), mites (N, Estimate = −0.361, sd = 0.176, p=0.40), and *E. uptoni* (OPG, Estimate = −3.53, sd = 1.51, p = 0.011), but increased infection abundance of *E. hungaryensis* (OPG, Estimate = 1.67, sd = 0.51, p = 0.0011). Infection dynamics also varied substantially across years (Figure 1; Figure 3; Table S3).

### Evidence of behaviour-mediated supplementation effects on parasites

We tested whether effects of supplementation on behaviour (Figure 2) were mediating those on the parasite community by adding behavioural metrics to the parasite models to test whether affected behaviours are influence infection. We found some support for the possibility that behavioural changes caused by supplementation may have subsequent effects on parasite infection. As shown, both network strength and sum of movement were significantly reduced by supplementation (Figure 2). Network strength in turn had a significant positive effect on *Eimeria hungaryensis* (Estimate = 1.19, sd = 0.32, p = 0.0002) and total *Eimeria* (Estimate = 0.99, sd = 0.31, p = 0.0012), and higher movement within the grids had a significant positive relationship with WMHV (Estimate = 0.70, sd = 0.25, p = 0.0042) and *Bartonella spp*. prevalence (Estimate = 0.45, sd = 0.18, p = 0.011) (Figure 3C).

### Effects of anthelmintic treatment on the parasite community

We investigated the impact of anthelmintic treatment on the parasite community, whereby treatment was administered randomly within grids for an additional experiment investigating the impact of treatment combined with nutrition on *H. polygyrus* (Sweeny *et al*. 2021c). As previously reported (Sweeny *et al*. 2021c), anthelmintic treatment reduced *H. polygyrus* infection abundance (Estimate = - 1.65, sd = 0.37, p < 0.0001). Removal of worms likewise influenced increased abundance of *Eimeria hungaryensis* infection (Estimate = 1.67, sd = 0.51, p = 0.0013), in line with previous results in wood mice demonstrating a negative interaction between these two parasites (Knowles *et al*. 2013). We found additional negative effects of anthelmintic treatment on mite infestation (Estimate = −0.36, sd = 0.18, p = 0.04) and wood mouse herpesvirus probability (Estimate = −1.30, sd = 0.44, p = 0.003).

## Discussion

Humans are exerting a growing influence on wildlife diet, leading to increased interest in the health effects of supplementation. A growing body of evidence has highlighted significant effects of diet and nutrition quality on disease in wild populations (Becker *et al*. 2015; Murray *et al*. 2016). Theoretical and empirical work show that outcomes vary widely depending on parasite traits and host responses (Becker *et al*. 2015; Becker & Hall 2014; Erazo *et al*. 2022, 2025). However, how such effects scale to multi-parasite systems within a single population remains unexplored. Here, we show that supplemental feeding altered the majority of parasites surveyed in a wild population of wood mice (*Apodemus sylvaticus*). Alongside changes to the parasite community, we found that supplementation drastically altered host behaviour and demography and had subtle yet consistent trends toward improved host condition and immunity.

Our findings demonstrate that the effects of supplementation can be diverse and hard to predict for co-infected hosts. Notably, the direction of response was not easily explained by parasite type (e.g. gastrointestinal, ectoparasite or bloodborne). We found that the majority of effects reduced infection intensity, yet increases also occurred, highlighting competing mechanisms. Among gastrointestinal helminths, the dominant nematode *H. polygyrus* declined under supplementation, consistent with improved host resistance and previous findings in this population from a smaller dataset (Sweeny *et al*. 2021c). In contrast, another nematode (*C. murissylvatici*) increased, demonstrating divergent outcomes even within similar parasite groups. Infections by the common gastrointestinal protozoan genus *Eimeria* were consistently reduced by supplementation across all species (with all but *E. apionodes* statistically significant). Such variation across parasite species suggests that resource supplementation influences the GI parasite community through multiple pathways rather than a single dominant mechanism. For example, because *C. murissylvatici* is very rare in this system (Figure 1), reduction of other members of the GI community (*H. polygyrus* and *Eimeria spp*.) induced by supplementation may directly decrease competition for space in the wood mouse gut, facilitating establishment of *C. murissylvatici*. These results support previous theoretical work (Erazo *et al*. 2025) which suggests the effects of resource supplementation among coinfecting parasites can be hard to predict, due to the balance of both direct and indirect responses to supplementation, mediated by coinfection interactions.

Supplementation effects were similarly mixed across ecto- and micro-parasites. Ticks (*Ixodes spp*.) were significantly reduced, while mite infestation increased. Supplementation increased *Bartonella spp*. and decreased wood mouse herpesvirus (WMHV), while fleas and *T. grosi* showed no detectable response. Meanwhile fleas and their vectored protozoan *T. grosi* show no response to supplementation, although fleas were very rare in this population (Figure 1). These contrasting responses reinforce that parasite life history, environmental transmission stages, and sensitivity to immune or behavioural changes can influence infection outcomes under resource supplementation.

Decreased infection across several parasite taxa and modest improvements in condition and immune response suggest that supplementation enhanced host resistance to at least part of the parasite community. High variation in condition effects observed here in the combined 2015-2017 dataset may suggest that natural food availability may modulate supplementation effects across years in this experiment, particularly as 2017 was a beech (*Fagus* spp.) mast year for these woodlands. Previous work from this population limited to 2015 and 2016 showed improvements to body mass and higher circulating *H. polygyrus*-specific IgG1 and total IgA following supplementation (Sweeny *et al*. 2021c), suggesting that supplementation effects on condition and immunity may be strongest when natural resources are a limiting factor (e.g., n non-mast years). It is possible that generally higher condition and immunity (despite variation in the effects) plays a role in reduction of infections with *Eimeria spp*., WMHV, and *Ixodes spp*. if supplementation increases resistance to multiple parasites.

The lack of consistent responses within transmission categories suggests that supplementation effects are shaped by a complex interplay of susceptibility and exposure processes (Erazo *et al*. 2022; Sweeny & Albery 2022). Host behaviour is intimately tied to pathogen exposure, and the widespread effects of supplementation on multiple aspects of behaviour observed here may play a role in transmission dynamics. For example, although *Eimeria spp*. infections in other systems are known to correlate negatively with body condition (Hakkarainen *et al*. 2007), they are also heavily influenced by exposure to infective oocysts in the environment, often driven by high burdens in juveniles (Chartier & Paraud 2012; Sweeny *et al*. 2022) and host density (Albery *et al*. 2025). We found that higher burdens of total *Eimeria* and *E. hungaryensis* infections were linked to more connected (‘strength’) and more mobile (‘sum movement’) individuals in this population. Supplementation reduced both social connections and movement, and therefore may have reduced exposure to *Eimeria* via these effects. Reduced movement and sociality may likewise have contributed to reductions of other parasites. Investigations in other *Apodemus* populations have found that males disproportionately contribute to transmission of WMHV (Erazo *et al*. 2021; Knowles *et al*. 2012), *H. polygyrus* (Ferrari *et al*. 2004), and other pathogens such as tick-borne encephalitis virus (Perkins *et al*. 2003; Rosà *et al*. 2019). In wood mouse populations, males tend to range the furthest (Attuquayefio *et al*. 1986); reduction in their home range (and social connections) due to supplementation may therefore reduce their contribution to transmission of *H. polygyrus* and WMHV, in line with observed decreases in both following supplementation. Although we did not identify definitive links between behaviour and *H. polygyrus* infections, host movement did have a positive effect on WMHV prevalence, and reduced movement among supplemented mice may therefore have reduced their WMHV prevalence.

The tick reduction observed here under supplementation may reflect a combination of increased host resistance and reduced exposure. Tick infestation was previously found to be higher on cattle consuming poor-quality feed (Sutherst *et al*. 1983) and on impalas in poor condition (Gallivan *et al*. 1995), and experimental manipulation of *Peromyscus* diet components increased resistance to ticks and capacity of hosts to cope with infestation (Blubaugh *et al*. 2023). However, tick transmission requires host contact with infectious stages in the environment, and therefore larger home ranges can increase exposure, as documented in sand lizards (Wieczorek *et al*. 2020). By contrast, mites often benefit from hosts in good condition (Dube *et al*. 2018), are directly transmitted between hosts, and synchronise reproduction with their hosts (Christe *et al*. 2000). Taken together, reductions in ticks may reflect both lower exposure due to reduced movement and improved nutritional status, whereas mite increases may be driven by a combination of host condition, higher local density, and increased reproductive investment in supplemented individuals. Further work following individuals for a longer timespan is required to resolve these relationships, as individuals were followed for only two weeks each in 2015-2016 and behavioural effects may occur with greater time lags. Finally, reductions in *Ixodes spp*. in this wood mouse population may also reflect host community-level responses to supplementation. Experimental field sites were also home to bank voles (*Clethrionomys glareolus*), and if supplementation increased reproduction and population size of alternative competent hosts, then increased availability of those hosts may have contributed to decreased infestation of ticks on wood mice (Ostfeld & Keesing 2000). Future work monitoring both multi-host and multi-parasite responses to changing resources could further resolve these mechanisms.

Our results support a growing body of evidence across host systems that show significant effects of diet and nutrition quality on disease in wild populations (Becker *et al*. 2015; Murray *et al*. 2016), and complement findings from recent work showing that supplemental feeding similarly reduced space use of wild bank voles (Mistrick *et al*. 2024). Although the bank vole study employed a similar factorial experimental design of supplementation and anthelmintic treatment, Mistrick et al. (2024) found that worm removal produced an opposite effect to supplementation by increasing space use; however, we did not observe significant effects of anthelmintic treatment on any metrics of behaviour in this study (Figure 3). This discrepancy may be due to several reasons. First, the observed increase in space use following anthelmintic treatment in the bank vole study emerged in the autumn following summer treatment, indicating that a substantial time lag may be required to observe treatment-induced behavioural changes. In contrast, individuals in our study were monitored for shorter time periods in two of three years, potentially missing delayed responses. Second, it has been proposed that increased movement following deworming is attributable to reduced sickness behaviours meant to conserve energy (Ghai *et al*. 2015). Because supplementation occurred before deworming in our study, treated individuals may have already had reduced sickness behaviours under increased resource availability, reducing detectable treatment effects. Finally, spatiotemporal context is crucial for interpreting infection dynamics (Sweeny *et al*. 2021b), and the populations in the two studies comprise very different species, locations, and years. As discussed above, one experimental year in the present study was a mast year, suggesting that these mice had generally higher baseline resources. Thus, helminth infections in our system may not have been sufficiently behaviourally costly across all years for treatment effects to manifest. The influence of spatiotemporal context on infection outcomes has been previously demonstrated in a single population of Rocky Mountain Elk, where supplemental feeding increased gastrointestinal helminths in winter but reduced them in spring (Hines *et al*. 2007). Extensions to this study quantifying availability of natural resources could provide further insights into how predictable host and parasite responses are to resource fluctuations across space and time.

Overall, this work demonstrated that resource supplementation drives divergent and taxon-specific responses across a diverse range of parasite species, even under a simple provisioning regime. These findings are relevant for understanding how anthropogenic change reshapes wildlife disease dynamics and how natural resource pulses may restructure parasite communities. Future work could use higher-resolution behavioural (e.g. proximity loggers (Kirkpatrick *et al*. 2021)) and immunological (e.g. RNA-seq (Watowich *et al*. 2024)) tools to test the mechanistic pathways linking susceptibility and exposure under resource change. Because parasite community effects are important for determining infection outcomes across scales, from the individual level (Telfer *et al*. 2010) up to disease emergence (Sweeny *et al*. 2021a), our study underscores that forecasting disease outcomes under environmental and anthropogenic change requires a community-level perspective.

## Supporting information

Electronic Supplementary Material

## Acknowledgements

We thank the Forestry Commission for permissions for fieldwork in Callendar Park. *Heligmosomoides polygyrus* excretory-secretory product (HES) used in immunological assays was supplied by Rick Maizels (University of Glasgow) and Amy Buck (University of Edinburgh). Many thanks also to many tireless assistants and volunteers involved in field and laboratory data collection.

## References

Albery, G.F., Sweeny, A.R., Corripio-Miyar, Y., Evans, M.J., Hayward, A.D., Pemberton, J.M., et al. (2025). Local and global density have distinct and parasite-dependent effects on infection in wild sheep. Parasitology, 152, 715–723.

Attuquayefio, D.K., Gorman, M.L. & Wolton, R.J. (1986). Home range sizes in the Wood mouse Apodemus sylvaticus: habitat, sex and seasonal differences. J. Zool., 210, 45–53.

Becker, D.J. & Hall, R.J. (2014). Too much of a good thing: resource provisioning alters infectious disease dynamics in wildlife. Biol. Lett., 10, 20140309–20140309.

Becker, D.J., Streicker, D.G. & Altizer, S. (2015). Linking anthropogenic resources to wildlife-pathogen dynamics: a review and meta-analysis. Ecol. Lett., 18, 483–495.

Blubaugh, C.K., Jones, C.R., Josefson, C., Scoles, G.A., Snyder, W.E. & Owen, J.P. (2023). Omnivore diet composition alters parasite resistance and host condition. J. Anim. Ecol., 92, 2175–2188.

Bogdziewicz, M. & Szymkowiak, J. (2016). Oak acorn crop and Google search volume predict Lyme disease risk in temperate Europe. Basic Appl. Ecol., 17, 300–307.

Calder, P.C. & Jackson, A.A. (2000). Undernutrition, infection and immune function. Nutr. Res. Rev., 13, 3–29.

Chartier, C. & Paraud, C. (2012). Coccidiosis due to Eimeria in sheep and goats, a review. Small Rumin. Res., 103, 84–92.

Christe, Arlettaz & Vogel. (2000). Variation in intensity of a parasitic mite (Spinturnix myoti) in relation to the reproductive cycle and immunocompetence of its bat host (Myotis myotis). Ecol. Lett., 3, 207–212.

Cox, F.E.G. (2001). Concomitant infections, parasites and immune responses. Parasitology, 122, S23–S38.

Cressler, C.E., Nelson, W.A., Day, T. & McCauley, E. (2014). Disentangling the interaction among host resources, the immune system and pathogens. Ecol. Lett., 17, 284–293.

Cross, P.C., Edwards, W.H., Scurlock, B.M., Maichak, E.J. & Rogerson, J.D. (2007). Effects of management and climate on elk brucellosis in the Greater Yellowstone Ecosystem. Ecol. Appl., 17, 957–964.

Dube, W.C., Hund, A.K., Turbek, S.P. & Safran, R.J. (2018). Microclimate and host body condition influence mite population growth in a wild bird-ectoparasite system. Int. J. Parasitol. Parasites Wildl., 7, 301–308.

Erazo, D., Pedersen, A.B. & Fenton, A. (2022). The predicted impact of resource provisioning on the epidemiological responses of different parasites. J. Anim. Ecol., 91, 1719–1730.

Erazo, D., Pedersen, A.B., Gallagher, K. & Fenton, A. (2021). Who acquires infection from whom? Estimating herpesvirus transmission rates between wild rodent host grousps. Epidemics, 35, 100451.

Erazo, D., Sweeny, A.,. Pedersen, A.B. & Fenton, A. (2025). Parasite responses to resource provisioning can be altered by within-host co-infection interactions. Proc. Biol. Sci.

Ferrari, N., Cattadori, I.M., Nespereira, J., Rizzoli, A. & Hudson, P.J. (2004). The role of host sex in parasite dynamics: field experiments on the yellow□necked mouse Apodemus flavicollis: Role of host sex in parasite dynamics. Ecol. Lett., 7, 88–94.

Gallivan, G.J., Culverwell, J., Girdwood, R. & Surgeoner, G.A. (1995). Ixodid ticks of impala (Aepyceros melampus) in Swaziland: effect of age class, sex, body condition and management. S. Afr. J. Zool., 30, 178–186.

Garnier, R., Cheung, C.K., Watt, K.A., Pilkington, J.G., Pemberton, J.M. & Graham, A.L. (2017). Joint associations of blood plasma proteins with overwinter survival of a large mammal. Ecol. Lett., 20, 175–183.

Ghai, R.R., Fugère, V., Chapman, C.A., Goldberg, T.L. & Davies, T.J. (2015). Sickness behaviour associated with non-lethal infections in wild primates. Proc. Biol. Sci., 282, 20151436.

Glaudas, X. & Alexander, G.J. (2017). Food supplementation affects the foraging ecology of a low-energy, ambush-foraging snake. Behav. Ecol. Sociobiol., 71.

Hakkarainen, H., Huhta, E., Koskela, E., Mappes, T., Soveri, T. & Suorsa, P. (2007). Eimeria-parasites are associated with a lowered mother’s and offspring’s body condition in island and mainland populations of the bank vole. Parasitology, 134, 23–31.

Hellard, E., Fouchet, D., Vavre, F. & Pontier, D. (2015). Parasite–Parasite Interactions in the Wild: How To Detect Them? Trends Parasitol., 31, 640–652.

Hines, A.M., Ezenwa, V.O., Cross, P. & Rogerson, J.D. (2007). Effects of supplemental feeding on gastrointestinal parasite infection in elk (Cervus elaphus): preliminary observations. Vet. Parasitol., 148, 350–355.

Kirkpatrick, L., Hererra Olivares, I., Massawe, A., Sabuni, C., Leirs, H., Berkvens, R., et al. (2021). ProxLogs: Miniaturised proximity loggers for monitoring association behaviour in small mammals. bioRxiv.

Knowles, S.C.L., Fenton, A. & Pedersen, A.B. (2012). Epidemiology and fitness effects of wood mouse herpesvirus in a natural host population. J. Gen. Virol., 93, 2447–2456.

Knowles, S.C.L., Fenton, A., Petchey, O.L., Jones, T.R., Barber, R. & Pedersen, A.B. (2013). Stability of within-host-parasite communities in a wild mammal system. Proc. Biol. Sci., 280, 20130598.

Mistrick, J., Veitch, J.S.M., Kitchen, S.M., Clague, S., Newman, B.C., Hall, R.J., et al. (2024). Effects of food supplementation and helminth removal on space use and spatial overlap in wild rodent populations. J. Anim. Ecol., 93, 743–754.

Moyers, S.C., Adelman, J.S., Farine, D.R., Thomason, C.A. & Hawley, D.M. (2018). Feeder density enhances house finch disease transmission in experimental epidemics. Philos. Trans. R. Soc. Lond. B Biol. Sci., 373, 20170090.

Murray, M.H., Becker, D.J., Hall, R.J. & Hernandez, S.M. (2016). Wildlife health and supplemental feeding: A review and management recommendations. Biol. Conserv., 204, 163–174.

Nagy, L.R. & Holmes, R.T. (2005). Food limits annual fecundity of a migratory songbird: An experimental study. Ecology, 86, 675–681.

Oro, D., Genovart, M., Tavecchia, G., Fowler, M.S. & Martínez-Abraín, A. (2013). Ecological and evolutionary implications of food subsidies from humans. Ecol. Lett., 16, 1501–1514.

Ostfeld, R.S. & Keesing, F. (2000). Biodiversity and disease risk: The case of Lyme disease. Conserv. Biol., 14, 722–728.

Pedersen, A.B. & Fenton, A. (2015). The role of antiparasite treatment experiments in assessing the impact of parasites on wildlife. Trends Parasitol., 31, 200–211.

Peig, J. & Green, A.J. (2009). New perspectives for estimating body condition from mass/length data: the scaled mass index as an alternative method. Oikos, 118, 1883–1891.

Perkins, S.E., Cattadori, I.M., Tagliapietra, V., Rizzoli, A.P. & Hudson, P.J. (2003). Empirical evidence for key hosts in persistence of a tick-borne disease. Int. J. Parasitol., 33, 909–917.

Petney, T.N. & Andrews, R.H. (1998). Multiparasite communities in animals and humans: frequency, structure and pathogenic significance. Int. J. Parasitol., 28, 377–393.

Randolph, S.E. & Storey, K. (1999). Impact of microclimate on immature tick-rodent host interactions (Acari: Ixodidae): implications for parasite transmission. J. Med. Entomol., 36, 741–748.

Rosà, R., Tagliapietra, V., Manica, M., Arnoldi, D., Hauffe, H.C., Rossi, C., et al. (2019). Changes in host densities and co-feeding pattern efficiently predict tick-borne encephalitis hazard in an endemic focus in northern Italy. Int. J. Parasitol., 49, 779–787.

Rue, H., Martino, S. & Chopin, N. (2009). Approximate Bayesian Inference for Latent Gaussian models by using Integrated Nested Laplace Approximations. J. R. Stat. Soc. Series B Stat. Methodol., 71, 319–392.

Sutherst, R.W., Kerr, J.D., Maywald, G.F. & Stegeman, D.A. (1983). The effect of season and nutrition on the resistance of cattle to the tick Boophilus microplus. Aust. J. Agric. Res., 34, 329.

Sweeny, A.R. & Albery, G.F. (2022). Exposure and susceptibility: The Twin Pillars of infection. Funct. Ecol., 36, 1713–1726.

Sweeny, A.R., Albery, G.F., Becker, D.J., Eskew, E.A. & Carlson, C.J. (2021a). Synzootics. J. Anim. Ecol., 90, 2744–2754.

Sweeny, A.R., Albery, G.F., Venkatesan, S., Fenton, A. & Pedersen, A.B. (2021b). Spatiotemporal variation in drivers of parasitism in a wild wood mouse population. Funct. Ecol., 35, 1277–1287.

Sweeny, A.R., Clerc, M. & Pontifes, P.A. (2021c). Supplemented nutrition decreases helminth burden and increases drug efficacy in a natural host–helminth system. of the Royal ….

Sweeny, A.R., Corripio-Miyar, Y., Bal, X., Hayward, A.D., Pilkington, J.G., McNeilly, T.N., et al. (2022). Longitudinal dynamics of co-infecting gastrointestinal parasites in a wild sheep population. Parasitology, 149, 1–12.

Telfer, S., Lambin, X., Birtles, R., Beldomenico, P., Burthe, S., Paterson, S., et al. (2010). Species interactions in a parasite community drive infection risk in a wildlife population. Science, 330, 243–246.

Ullman-Cullere, M.H. & Foltz, C.J. (2011). Body Condition Scoring: A Rapid and Accurate Method for Assessing Health Status in Mice, 1–5.

Watowich, M.M., Costa, C.E., Chiou, K.L., Goldman, E.A., Petersen, R.M., Patterson, S., et al. (2024). Immune gene regulation is associated with age and environmental adversity in a nonhuman primate. Mol. Ecol., 33, e17445.

Wieczorek, M., Rektor, R., Najbar, B. & Morelli, F. (2020). Tick parasitism is associated with home range area in the sand lizard, Lacerta agilis. Amphib-reptil.

