## Supplementary material for "Resource supplementation alters host exposure, susceptibility, and infection dynamics across a diverse parasite community": Electronic Supplementary Material

### S1 Methods

##### PCR protocols for blood-borne parasites

DNA was extracted from blood samples using a MagMAXTM-96 DNA Multi-Sample Kit (ThermoFisher Scientific) on a KingFisherTM Flex Purification System optimised for small blood volumes and mouse samples according to manufacturer instructions. This kit is designed for high throughput purification of DNA from animal tissue and blood and uses magnetic bead-based isolation and extractions were carried out according to manufacturer’s instructions. Briefly, 10 μL of dH2O was added to whole blood pellets and homogenised. Samples were lysed with proteinase K followed by a treatment with guanidinium thiocyanate-based solution. Following lysis, samples were mixed with isopropanol and combined with paramagnetic beads with a nucleic acid binding surface. The beads, with bound nucleic acid, are immobilized on magnets and washed to remove proteins and other contaminants. A second wash solution is used to remove residual binding solution and then the nucleic acid is eluted using a low-salt buffer.

Trypanosomes were detected using a nested PCR targeting a 530bp section of the 18S rRNA gene (Noyes *et al.* 2002). Primers TRY927F 5’-GAAACAAGAAACACGGGAG and TRY927R 5’-CTACTGGGCAGCTTGGA were used in the first round, and SSU561F 5’-TGGGATAACAAAGGAGCA and SSU561R 5’- CTGAGACTGTAACCTCAAAGC in the second round. Each reaction contained 2μl genomic DNA, 0.1mM each dNTP, 0.2μM each primer, 0.8mM MgCl2, 0.5U Platinum Taq DNA Polymerase (Invitrogen) with the accompanying buffer at 1x concentration. A touchdown PCR profile was used, with initial denaturation at 96°C for 3min, followed by 20 cycles of 96°C for 30s, 60°C for 1min (decreasing by 0.5°C each cycle until 55°C), and 72°C for 90s, followed by a final extension at 72°C for 10min. In the event that any negative control was positive, the entire PCR plate was repeated. 5μl of all PCR products were run on 2% agarose gels stained with ethidium bromide and visualized under UV light. For the trypanosome PCR, samples showing an approx. 500bp band were scored as positive. All positive samples were sequenced and across the 502bp sequence obtained (excluding primers) all were 100% identical to Trypanosoma grosi (Genbank accession: AB175624).

Wood Mouse Herpes Virus (WMHV) was detected via nested PCR using pan-herpesvirus primers targeting a 160 bp region of the DPOL gene: ILK+ 5′-ATAAACAACAGCTGGCCATCAA-3′ and KG1+ 5′- CTGACCAGATCCACCCCTTT-3′ in the first round, followed by TGV+ 5′- TGTAATTCTGTCTATGGCTTCACAGGAGT-3′ and IGY+ 5′- AAGAGAATCTGTGTCTCCATAAAT-3′ in the second round (Ehlers *et al.* 2008). Reactions were run in 25 μl volumes, containing 0.4 μM each primer (Metabion), 0.2 mM dNTPs, 3 mM MgCl2, one unit GoTaq DNA Polymerase (Promega), 5 μl 5× GoTaq Flexi Buffer (Promega) and 2 μl template DNA. The following PCR conditions were used for either 25 cycles (first round) or 35 cycles (second round): initial denaturation at 95 °C for 3 min, cycles of 95 °C for 20 s, 61 °C for 30 s and 72 °C for 30 s, and a final extension of 72 °C for 10 mins.

*Bartonella* spp. were detected via genus-specific , semi-nested PCR targeting a fragment of 16S-23S internal transcribed spacer region, using previously described methods (Telfer *et al.* 2005) with several modifications. 8.8µl H2O, 2.16µl 25mM MgCl2, 0.27µl each of primer bigF (5’-TTG ATA AGC GTG AGG TC ) and bogR (5’-TGC AAA GCA GGT GCT CTC CCA) and 2 µl of template DNA were used for the first round of PCR. In the second round of PCR, 9.8µl H2O, 2.16µl 25mM MgCl2, 0.27µl each of 10µM primer bigF (5’-TTG ATA AGC GTG AGG TC ) and bigR (5’-TCC CAG CTG AGC TAC G) and 1µl of first round’s PCR product was used as the template. The PCR was amplified with the following conditions for round 1: 96°C for 3min, 96°C for 10s, 61°C for 10sec and then decrease by 0.5°C every cycle, 72°C for 50s, back to step 2 for 12 more times, 96°C for 10s, 55°C for 10s, 72°C for 50s, back to 6 for 7 more times, and 8°C forever. Round 2 conditions were as follows: 96°C for 3min, 96°C for 10s, 61°C for 10sec and then decrease by 0.5°C every cycle, 72°C for 50s, back to step 2 for 12 more times, 96°C for 10s, 55°C for 10s, 72°C for 50s, back to 6 for 21 more times, and 8°C forever.

##### Immune and condition assays

We used Enzyme-Linked Immunosorbence Assays (ELISAs) to measure (1) total faecal IgA concentration and (2) serum *H. polygyrus*-specific IgG1 antibody titres for each mouse at each capture/sampling point as previously described (Sweeny *et al.* 2021). We calculated total faecal IgA concentration by extrapolation from a standard curve of known concentrations from a synthetically manufactured standard antibody. *H. polygyrus*-specific IgG1 was calculated as a relative concentration to a positive reference sample consisting of sera from *Mus musculus* experimentally infected with H. polygyrus in the laboratory. Plates were prepared with serial dilutions of reference and experimental samples, and a dilution factor of 1:200 was selected for calculation of relative antibody concentrations. Standardised IgG1 concentrations were calculated by plate as follows: (Sample OD1:200- Mean Blanks)/ (Positive reference OD1:200-Mean Blanks). We assigned a value of 0 to samples for which the OD did not exceed 3x SD of control blanks. We refer to both IgA and IgG1 values as ‘antibody concentration’.

Nutritional status was assessed by quantifying circulating serum albumin concentration from the serum samples. Serum albumin is a dynamic and long-lived plasma protein which has important functions for multiple physiological roles (Garnier *et al.* 2017). Assays were optimized from (Garnier *et al.* 2017) for use on mouse samples using samples from our University of Edinburgh colony of formerly- wild wood mice. Samples for the serum albumin assays were diluted 1:4 in Mill-Q water and 5uL was added to each well and adjusted to a total volume of 50uL with albumin buffer. 100uL of bromocresol green reagent (prepared according to kit guidelines, Biovision Albumin (BCG) Colorimetric Assay Kit) was added to each well. Plates were shaken for 45”, incubated at RT for 20’ and read at 650nm. Samples for total protein assays were diluted 1:60 in Milli-Q water and 10uL of sample was added to each well. 300uL of Coomassie Reagent (Thermoscientific). Coomassie Plus Bradford Assay) was added to each well. Plates were shaken for 45”, incubated at RT for 10’ and read at 570nm. Samples were run in duplicate for both assays. Two standard dilutions of known concentrations of BSA were included for each plate in both assays. Standard curves were fit using a 4-parameter logistic regression and sample concentrations were determined by plate using the standard curves. Concentrations were corrected by sample dilution factors and expressed as ug/uL for analyses.

### S2 Results

#### Table captions

Table S1. Estimates, standard deviation, 95% credibility intervals, and symmetric Kullback-Leibler divergence of posterior effects for all response variables and effects for our behaviour model set.

Table S2. Estimates, standard deviation, 95% credibility intervals, and symmetric Kullback-Leibler divergence of posterior effects for all response variables and effects for our immunity and condition model set.

Table S3. Estimates, standard deviation, 95% credibility intervals, and symmetric Kullback-Leibler divergence of posterior effects for all response variables and effects for our parasite model set.

### Supplementary material references

Ehlers, B., Dural, G., Yasmum, N., Lembo, T., de Thoisy, B., Ryser-Degiorgis, M.-P., *et al.* (2008). Novel mammalian herpesviruses and lineages within the *Gammaherpesvirinae* : Cospeciation and interspecies transfer. *J. Virol.*, 82, 3509–3516.

Garnier, R., Cheung, C.K., Watt, K.A., Pilkington, J.G., Pemberton, J.M. & Graham, A.L. (2017). Joint associations of blood plasma proteins with overwinter survival of a large mammal. *Ecol. Lett.*, 20, 175–183.

Noyes, H.A., Ambrose, P., Barker, F., Begon, M., Bennet, M., Bown, K.J., *et al.* (2002). Host specificity of Trypanosoma (Herpetosoma) species: evidence that bank voles (Clethrionomys glareolus) carry only one T. (H.) evotomys 18S rRNA genotype but wood mice (Apodemus sylvaticus) carry at least two polyphyletic parasites. *Parasitology*, 124, 185–190.

Sweeny, A.R., Clerc, M. & Pontifes, P.A. (2021). Supplemented nutrition decreases helminth burden and increases drug efficacy in a natural host–helminth system. *of the Royal …*.

Telfer, S., Bown, K., Sekules, R., Begon, M., Hayden, T. & Birtles, R. (2005). Disruption of a host-parasite system following the introduction of an exotic host species. *Parasitology*, 130, 661–668.
